# Who am I? Optimal Tissue for Germline Genetic Testing Post-Stem Cell Transplantation

**DOI:** 10.1101/2025.01.20.630521

**Authors:** Matthias Mertens, Mona Sadlo, Jörn-Sven Kühl, Klaus Metzeler, Louisa Zschenderlein, Jeanett Edelmann, Claudia Lehmann, Sarah Thull, Mert Karakaya, Clara Velmans, Theresa Tumewu, Matthias Böhme, Christina Klötzer, Anne Weigert, Vladan Vucinic, Julia Hentschel, Mareike Mertens

## Abstract

With advancements in genetic diagnostics and genotype-based therapeutics, the demand for germline genetic testing in post-hematopoietic stem cell transplantation patients is increasing. Due to genetic chimerism, blood samples can no longer be used for germline testing after transplantation. This study aims to identify the most suitable tissue for germline analysis following stem cell transplantation by investigating alternative tissue sources. Buccal swab, eyebrow hair, and nail samples were analyzed for donor-derived DNA using next-generation sequencing and short tandem repeat analysis, with linear regression used for evaluation. Factors such as HLA match, transplantation type, sex, and time after transplantation were also evaluated for their effect on donor-derived DNA share. Buccal swab and nail samples exhibited 25% and 22% higher proportions of donor-derived DNA compared to eyebrow hair follicles, respectively. The median donor DNA share in eyebrow hair follicles was 1% for NGS and 3% for STR. Factors such as matched related donors, higher HLA match, different donor-recipient sex, and longer time post-transplantation correlated with lower donor DNA shares. Eyebrow hair follicles are a promising tissue for accurate germline genetic testing in post-SCT patients. Patient characteristics like donor relatedness, HLA match, sex match, and time after transplantation should be considered.

## I. Introduction

The integration of genetic information into clinical decision-making has revolutionized personalized medicine, allowing for the development of more precise treatment strategies. Therapies tailored to individuals with inherited pathogenic variants, such as BRCA1 and BRCA2 in familial breast and ovarian cancer, pathogenic RPE65 variants in Leber congenital amaurosis, or LDLR pathogenic variants in familial hypercholesterolemia, depend on precise genetic profiling to optimize therapeutic efficacy and mitigate adverse outcomes, to name some of them (1–3). Additionally, patients with conditions like Lynch syndrome or long QT syndrome require specific genetic testing to guide their treatment and prevent life-threatening complications (3–5). Moreover, pharmacogenetic interventions necessitate preemptive germline testing to identify polymorphisms that influence drug metabolism, as seen with DPYD variants in patients undergoing fluoropyrimidine therapy (6).

Stem cell transplantation plays a pivotal role in the treatment of various diseases, especially in hematological neoplasia, by restoring bone marrow function and providing graft-vs-leukemia effect. Therefor routine blood-based genetic testing methods fail to account for the presence of recipient-derived DNA in transplanted individuals. Several studies asserted that hair was devoid of donor-derived DNA (7). Jacewicz et al. have presented evidence to the contrary, demonstrating the presence of donor-derived DNA in hair tissue (8). It has also been observed that donor-derived DNA can be detected in tissues other than blood or hair cells, such as nails and buccal swab material, following stem cell transplantation (7,9–19). The genetic chimerism in those tissues limits the reliability and accuracy of germline testing in this patient population. These findings prompt the need to explore the most suitable tissue for genetic testing, particularly with the advent of new technologies like next-generation sequencing, long read sequencing and RNA Seq.

To determine the optimal approach for detecting donor-derived DNA, we compare next-generation sequencing (NGS) and short tandem repeat (STR) analysis. We investigated tissues that can potentially overcome the limitations posed by genetic chimerism. These include buccal swab samples, nail samples, and hair samples, each offering unique advantages for germline genetic testing. All these tissues provide a non-invasive to minimal invasive and easily accessible source of genetic material. Our goal is to identify the tissue that consistently exhibits the lowest donor-to-recipient ratio across all patients, which is crucial for accurate genetic diagnosis of the recipient. Additionally, we explore patient characteristics that may correlate with either a particularly high or particularly low donor percentage, providing further insights into the dynamics of chimerism in post-stem cell transplantation individuals.

## II. Materials and Methods

### Cohort

The study includes 46 participants who underwent stem cell transplantation, with 33 individuals younger than 18 years and 13 individuals being 18 years or older. 45 of these patients underwent allogeneic stem cell transplantation, while one patient received an autologous stem cell transplant, serving as an internal control. The predominant diagnosis among the participants was leukemia, affecting 19 out of 46 individuals. Other conditions included lymphoma (2 patients), x-linked adrenoleukodystrophy (5 patients), myelodysplastic neoplasia (4 patients), hemophagocytic lymphohistiocytosis (3 patients), and combined immunodeficiency (3 patients). No proband in our study experienced a recurrence of the underlying dissease, as evidenced by both absence of recurrence in blood samples and documentation in clinical reports.

In total, 30 evaluable sets were obtained, including all tissues such as nails, buccal swabs, and eyebrow hair follicles. Written consent was obtained from all participants or their legal guardians. The study protocol underwent thorough review and received approval from the ethics commission of the Medical Faculty of the University of Leipzig (186/21-ek). Additionally, the study is registered in a national register (drks.de, identifier: DRKS00032352).

### Material

We analyzed three materials: buccal swab, nail, and eyebrow hair samples. All these tissues offer a non-invasive and easily accessible DNA source. We collected the tissue samples during the transplant follow-up. To attribute individual gene sequences to either the donor or recipient, we established genetic profiles using donor DNA reference samples from pre-transplantation blood samples.

The buccal swab samples were obtained using the ORAcollect-DNA kit (DNA Genotek Inc, Stittsville, Canada), following the recommended collection technique. To ensure an adequate amount of DNA in our nail samples, we required at least five fingernail or toenail samples collected after a 2-week period without cutting nails. For eyebrow hairs, we required 15 hairs with hair roots. The buccal swab, nails, and eyebrow hairs were stored at room temperature until DNA isolation.

## Methods

### DNA Extraction

We extracted DNA from buccal swabs using the MagCore Genomic DNA Whole Blood Kit 101 and the MagCore® HF 16 Plus II instrument (RBC Bioscience, New Taipei City, Taiwan). For nail DNA extraction, we used the DNeasy Blood & Tissue Kit (Qiagen, Hilden, Germany). For eyebrow DNA extraction, we utilized the innuPREP Forensic Kit (Analytic Jena, Jena, Germany). Furthermore, we use DNA from blood samples collected before transplantation as reference materials for donors and recipients. We measured DNA concentration using the Tecan Plate Reader Infinite® 200 PRO (Tecan, Männedorf, Switzerland). In cases of low DNA concentrations, we employed NanoDrop 2000 (Thermo Fisher Scientific, Massachusetts, USA) or Qubit 4.0 (Thermo Fisher Scientific, Waltham, Massachusetts, USA). Our DNA sample analysis employed two distinct approaches: the NGS-RC-PCR-based SNP assay and a short tandem repeat (STR) assay.

### Next generation sequencing

We conducted comprehensive DNA analysis on buccal swab, nail, and hair samples using an NGS-RC-PCR-based SNP assay. This assay involved 34 loci, allowing us to discern and quantify the presence of donor-derived DNA in each tissue type (see Appendix Table A.2). The analysis utilized the EasySeq™ NGS Reverse Complement PCR Human Exome Sample Tracking and Identification Kit from NimaGen (Nijmegen, Netherlands). NGS analysis was conducted using either NextSeq 550 or NovaSeq 6000 from Illumina (San Diego, California, USA). NGS sequence analysis was facilitated by the Varvis® software from Limbus (Rostock, Germany).

### Short tandem repeat (STR) assay

The STR assay encompassed the examination of 27 STR markers, including the Amelogenin locus, through an in-house assay (see Appendix Table A.1). For PCR amplification, we utilized the Multiplex PCR Kit (Qiagen, Hilden, Germany), necessitating a pre-diluted DNA concentration of 40 ng/µl. Subsequently, the PCR products underwent separation via capillary electrophoresis using the ABI 3500 Genetic Analyzer (Thermo Fisher Scientific, Waltham, Massachusetts, USA). The fragment sizes were analyzed through the GeneMapper™ Software 5 (Thermo Fisher Scientific, Waltham, Massachusetts, USA).

### Calculating donor-derived DNA-shares

In our empirical analysis, we utilized the statistical software Stata (StataCorp LLC, College Station, Texas, USA). We developed a program code to categorize the results of both STR and NGS analyses for each marker as either “informative” or “not informative.” By utilizing our code to identify uninformative patient-marker combinations, we minimized potential human reporting errors. We further computed the donor DNA share for each patient-marker combination based on the measured genetic markers. For NGS, a patient-marker combination was deemed not informative in the following cases: i) insufficient coverage from NGS, ii) shared base between donor and recipient, iii) allelic fractions not totaling 100%, allowing for a permissible deviation of +/-3% (values were rescaled to 100%), or iv) identification of technical artifacts or mosaicism resulting in hybrid base combinations beyond the anticipated scope of donors or recipients.^ii^

In the context of STR analysis, we classify patient-marker as not informative in the following cases: i) the absence of a sufficiently strong signal, leading to missing information, ii) the presence of identical microsatellite profiles between the donor and recipient, and iii) discrepancies between the marker-specific microsatellite profile of donor or recipient and the observed allele length in the donor or recipient derived samples, which can result from measurement errors or mosaic variants. To maintain consistency across analyses, we excluded spots at Y-chromosome and Amelogenin markers in NGS and STR analysis, but we utilized this information to ensure the correct sample assignment.

Out of a total of 3,689 patient-marker combinations for NGS and 3,120 for STR, we classified 2,140 and 1,680, respectively as informative. To address potential outliers, our statistical analysis relies on patient-specific median values derived from the measured donor DNA shares.

To validate the precision of our methods, we conducted a dilution series analysis using diluted DNA from two control individuals that did not underwent stem-cell transplantation. Figure 1 illustrate the results of this analysis for NGS and STR. In both Panels the x-axes represent the true DNA shares of eight diluted samples, ranging from 2 % to 50 %. The y-axes display the measured DNA proportions based on our NGS/STR analysis. The light shaded areas represent the density of the distribution of DNA proportions within each dilution across all markers. In addition, the panels show boxplots for each dilution where median values are marked with a white dot.

**FIGURE 1.:**
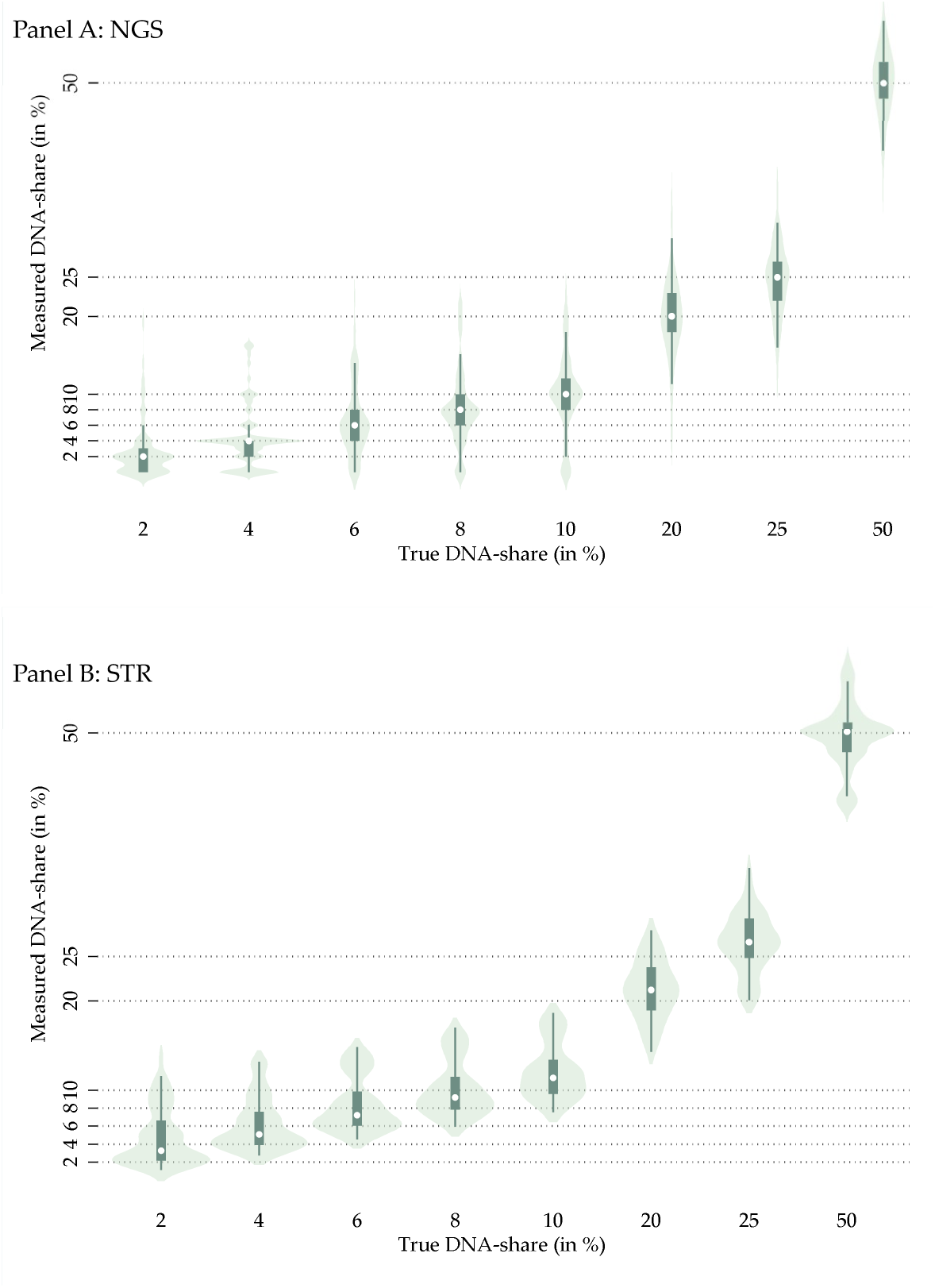
Dilution series NGS and STR.

Remarkably, our calculated median values closely match the true DNA proportions, providing strong evidence for the reliability of our methodology. Some individual marker values exhibit slight deviations from the true DNA proportions, likely reflecting measurement, and human error during laboratory procedures. However, these discrepancies remain within acceptable limits.

### Empirical analysis

#### Data collection and Overview

Table 1 provides a comprehensive overview of our final dataset, reporting summary statistics for 45 patients in our cohort. Our dataset includes information on donor DNA-shares, DNA concentration used in both Next-Generation Sequencing (NGS) and Short Tandem Repeat (STR) analysis, as well as quality parameters such as the amount of reads and average depth per tissue for NGS. These parameters are crucial for our quantitative analysis as they can potentially impact the measured donor-derived DNA-shares. Nails exhibited the highest values regarding reads and depth, with an average of 180,013 total reads per sample and an average depth of 1,507 per sample. In contrast, eyebrow hair follicles demonstrated lower total reads (70,578 per sample) and a lower average depth (639 per sample). Additionally, Table 1 reports a range of cohort characteristics that will serve as control variables.

**Table 1.** Summary statistics.

| SUMMARY STATISTICS |  |  |  |  |  |  |
| --- | --- | --- | --- | --- | --- | --- |
| Variable | Mean<br>(1) | Median<br>(2) | SD<br>(3) | Max<br>(4) | Min<br>(5) | N<br>(6) |
| DNA-concentration, nail sample (ng/μl) | 39.37 | 23.40 | 43.92 | 206.60 | 1.90 | 45 |
| DNA-concentration, buccal swab sample (μ/l) | 21.20 | 15.80 | 19.41 | 92.50 | 2.80 | 45 |
| DNA-concentration, eyebrow follicle sample (in μ/l) | 10.62 | 7.90 | 7.99 | 29.50 | 2.80 | 28 |
| Total reads per nail sample (NGS) | 180,013 | 91,185 | 347,303 | 2,215,796 | 1,935 | 45 |
| Average depth per nail sample (NGS) | 1,507 | 762 | 2,851 | 17,270 | 1 | 45 |
| Total reads per buccal swab sample (NGS) | 569,727 | 101,578 | 1,482,128 | 6,446,105 | 6,148 | 45 |
| Average depth per buccal swab sample (NGS) | 5,466 | 816 | 14,862 | 64,418 | 104 | 45 |
| Total reads per eyebrow follicle sample (NGS) | 70,578 | 70,739 | 55,794 | 195,410 | 3,853 | 28 |
| Average depth per eyebrow follicle sample (NGS) | 639 | 607 | 466 | 1,634 | 35 | 28 |
| Age at transplantation (years) | 18.77 | 8.00 | 22.30 | 72 | 0.40 | 45 |
| Days after transplantation | 1,319 | 965 | 1,319 | 6,339 | 97 | 45 |
| Gender donor (1=male/0=female) | 0.78 | 1.00 | 0.42 | 1.00 | 0.00 | 45 |
| Gender recipient (1=male/0=female) | 0.58 | 1.00 | 0.50 | 1.00 | 0.00 | 45 |
| Transplant type (1=bone marrow/0=periph. stem cell) | 0.51 | 1.00 | 0.51 | 1.00 | 0.00 | 43 |
| Complication after transplantation (1/0) | 0.53 | 0.00 | 0.49 | 1.00 | 0.00 | 43 |
| Graft vs. Host Disease (1/0) | 0.60 | 1.00 | 0.49 | 1.00 | 0.00 | 43 |
| HLA-match (scale from 1-10, 10 being best) | 9.9 | 10.00 | 0.29 | 10.00 | 8.00 | 44 |
| Blood group compatibility (1/0) | 0.49 | 0.00 | 0.51 | 1.00 | 0.00 | 43 |
| Matched related donor (1/0) | 0.24 | 0.00 | 0.43 | 1.00 | 0.00 | 45 |
| Resus factor match (1/0) | 0.73 | 1.00 | 0.45 | 1.00 | 0.00 | 45 |

Table 1 presents summary statistics of our data. Columns 1, 2, 3, 4, 5, and 6 report means, medians, standard deviations maxima, minima, and the number of individuals with non-missing information. The data has been self-collected as described in Materials and Methods.

#### Regression Analysis

To investigate the statistical variations in donor-derived DNA among different tissue types, we use a linear regression analysis. We estimate the following equation using ordinary least squares (OLS):

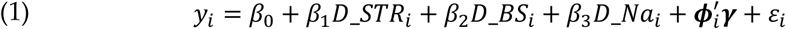

where, *y*_*i*_ denotes the donor DNA-share of patient *i*. *D_STR_i_, D_BS_i_, D_Na_i_* are dummies for data points from STR, buccal swab tissues, and nail tissues. The omitted categories are NGS and eyebrow hair follicle. The coefficient β thus indicates the difference in donor DNA shares between STR and NGS analyses, whereas β_n_ and β measure the difference in donor DNA between buccal swab and eyebrow hair follicle tissues and between nail and eyebrow hair follicle tissue. These are also our coefficients of interest. β captures the intercept, the vector φ^’^_*i*_ contains a series of control variables with coefficient values γ. The term ε represents the statistical error term. Given that individuals may enter multiple times with multiple tissue samples into the regression analysis, we cluster standard errors at the patient-level.

## III. Results

### Tissue-Specific Differences in Donor DNA-Shares

Figure 2 reports donor derived DNA-shares for each analyzed tissue. The Figure shows box plots of patient-specific median DNA-shares for each tissue and method.^iii^ We find pronounced differences in donor derived DNA-shares across the distinct tissue types. Notably, nail and buccal swab tissues exhibit high levels of donor derived DNA with median values of 8 % (NGS) and 16 % (STR) for nails, and 20 % (NGS) and 24 % (STR) for buccal swab tissues, while eyebrow hair follicle samples display considerably lower levels with median levels of 1 % (NGS) and 3 % (STR).

**FIGURE 2:**
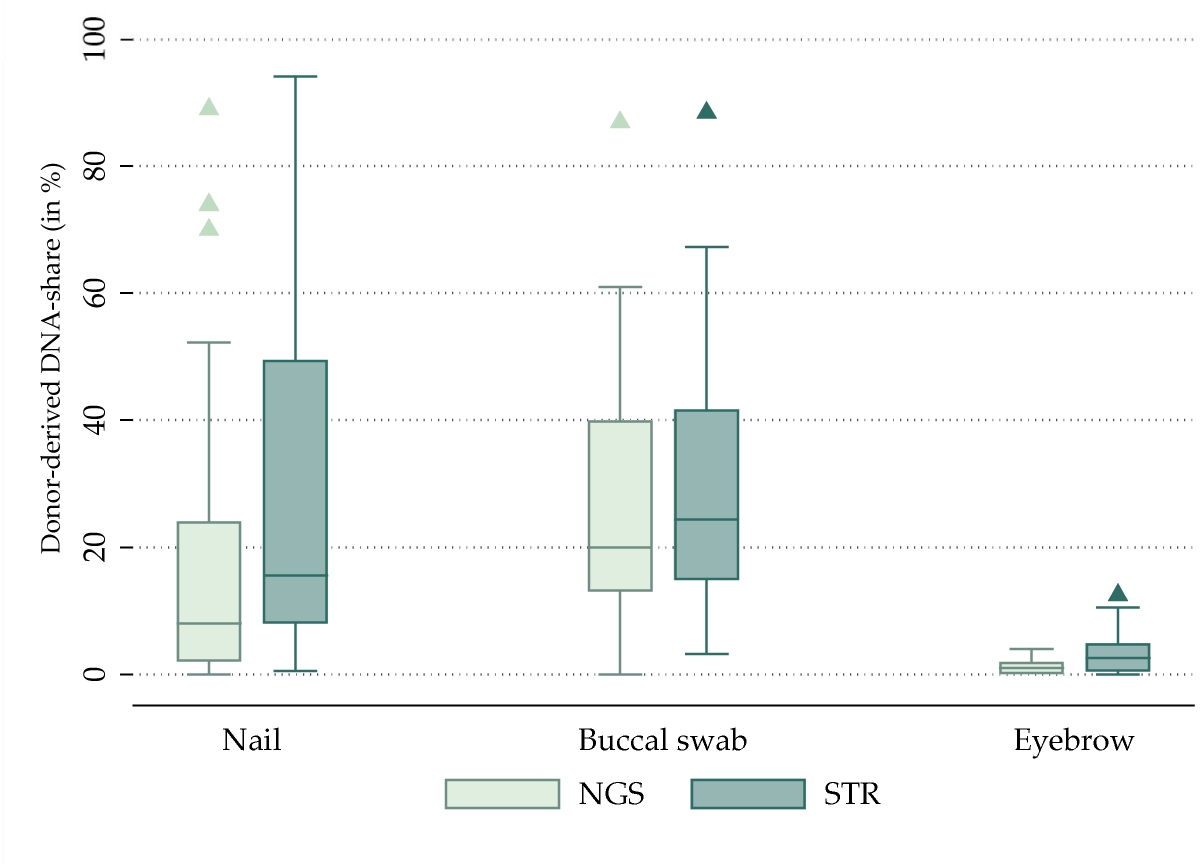
Differences in donor-derived DNA-shares across tissues.

### Regression Analysis Results – Key Findings

Table 2 presents the results obtained from estimating various specifications of regression equation (1) using ordinary least squares regression. Columns 1 to 4 display the results obtained when pooling NGS and STR datapoints. Columns 5 and 6 present separate regressions for STR and NGS data.

**Table 2.** Regression analysis.

| REGRESSION ANALYSIS |  |  |  |  |  |  |
| --- | --- | --- | --- | --- | --- | --- |
| *Dep. Variable: Median donor share* |  |  |  |  |  |  |
|  | NGS & STR | NGS & STR | NGS & STR | NGS & STR | STR | NGS |
|  | (1) | (2) | (3) | (4) | (5) | (6) |
| STR (1/0) | 0.061***<br>(0.007) | 0.062***<br>(0.007) | 0.062***<br>(0.007) | 0.059***<br>(0.007) |  |  |
| Buccal swab (1/0) |  | 0.251***<br>(0.027) | 0.252***<br>(0.036) | 0.227***<br>(0.032) | 0.240***<br>(0.036) | 0.225***<br>(0.035) |
| Nail (1/0) |  | 0.220***<br>(0.038) | 0.221***<br>(0.038) | 0.220***<br>(0.040) | 0.261***<br>(0.047) | 0.186***<br>(0.037) |
| Log (DNA-concentration) |  |  |  | -0.006<br>(0.024) | -0.006<br>(0.028) | 0.002<br>(0.023) |
| Same sex (1/0) |  |  |  | 0.078***<br>(0.026) | 0.082***<br>(0.030) | 0.071**<br>(0.027) |
| Log (age at transplantation) |  |  |  | 0.031*<br>(0.017) | 0.035*<br>(0.019) | 0.027<br>(0.017) |
| Log (days after transplantation) |  |  |  | -0.042**<br>(0.018) | -0.042**<br>(0.019) | -0.035*<br>(0.019) |
| Bone marrow transplant (1/0) |  |  |  | 0.040<br>(0.039) | 0.053<br>(0.043) | 0.018<br>(0.042) |
| Complication after transplantation (1/0) |  |  |  | -0.008<br>(0.034) | -0.003<br>(0.039) | -0.009<br>(0.033) |
| Blood group compatibility (1/0) |  |  |  | 0.026<br>(0.031) | 0.031<br>(0.035) | 0.021<br>(0.036) |
| HLA match 10 (1/0) |  |  |  | -0.186***<br>(0.068) | -0.196***<br>(0.072) | -0.153*<br>(0.082) |
| Graft vs. Host Disease (1/0) |  |  |  | 0.022<br>(0.036) | 0.023<br>(0.042) | 0.017<br>(0.033) |
| Matched rhesus factor (1/0) |  |  |  | 0.032<br>(0.036) | 0.034<br>(0.041) | 0.030<br>(0.034) |
| Matched related donor (1/0) |  |  |  | -0.121***<br>(0.036) | -0.136***<br>(0.042) | -0.106***<br>(0.035) |
| Log (reads) |  |  |  |  |  | -0.029<br>(0.020) |
| Log (depth) |  |  |  |  |  | 0.017<br>(0.015) |
| Regression constant | 0.171***<br>(0.018) | -0.009**<br>(0.004) | -0.009<br>(0.024) | 0.043<br>(0.096) | 0.070<br>(0.105) | 0.246*<br>(0.143) |
| Patient fixed effects | no | no | yes | no | no | no |
| Observations | 235 | 235 | 235 | 217 | 108 | 109 |
| R-squared | 0.019 | 0.227 | 0.581 | 0.374 | 0.381 | 0.362 |

Table 2 reports results from estimating equation (1) for separate samples using ordinary least squares. Columns 1-4 display results from pooling NGS and STR datapoints, while columns 5-6 report results from separate regressions for STR and NGS datapoints. Standard errors are clustered at the patient level and reported in parentheses. Significance: \*10 percent, \*\*5 percent, \*\*\*1 percent.

Utilizing the STR approach consistently resulted in on average 6 percent points higher donor-derived DNA shares compared to NGS. This effect is statistically significant at the 1% level (column 1).

The most prominent result in our study pertains to tissue-specific differences. Eyebrow follicles show by far the lowest amount of donor derived DNA. Buccal swab tissues consistently exhibited on average 25 percentage points higher donor DNA shares than eyebrow follicle tissues, with nail tissues also displaying an on average of 22 percentage points higher donor derived DNA shares. These differences are consistently observed across both NGS and STR assays and are statistically significant at the 1 % level (column 2). In column 3, we run the same regressions but control for patient fixed effects, such that we compare differences in donor-derived DNA shares across tissues within patients only. Our results are almost unchanged. As shown in column 4, incorporating control variables, such as DNA concentration and various cohort characteristics, does not alter our primary findings. Importantly, the influence of DNA concentration on donor derived DNA shares is small and statistically insignificant. When analyzing NGS and STR data separately, our key results also remain robust (columns 5 and 6).

### Regression Analysis Results – Other Clinically Relevant Results

Beyond tissue-specific disparities, our analysis reveals further novel insights. Donor and recipient sex-matching correlated with an approximately 7-8 percentage points higher donor derived DNA shares (columns 4-6 in Table 2). Additionally, a longer time after transplantation is linked to lower donor derived DNA shares. Notably, a better HLA-match is related to lower donor DNA shares. The effect is large and amounts to 15-20 percentage points, depending on the specification. Finally, we find strong evidence that stem cell transplantation from a matched related donor correlates with a lower donor DNA-share of 12-14 percentage points, depending on the specification. Together with the results on the HLA-match, this suggests that a better match quality has a strong negative impact on the donor DNA-share in the examined materials of the recipient.

## IV. Discussion

Our study aimed to identify the most suitable tissue for germline genetic testing using up-to-date genetic testing methods such as next generation sequencing after stem cell transplantation. The primary focus was to identify the tissue exhibiting the least donor-derived DNA share. Some studies already detected donor derived DNA in nail samples, buccal swab samples, and hair samples using a small amount of probands, not using eyebrow hair follicle samples (7–17). Our study features the largest patient sample to date and is the first to employ Next-Generation Sequencing (NGS) on buccal swab samples, nail samples, and eyebrow hair follicle samples for detecting donor-derived DNA. Moreover, we contribute by relating various patient characteristics to observed donor-derived DNA shares.

We show that eyebrow hair follicle samples exhibited an exceptionally low percentage of donor-derived DNA, with only 1 % in the NGS assay on average. Using regression analysis, we show that buccal swab and nail tissues are on average characterized by an at least 20 percentage point higher donor derived DNA share. This result is highly statistically significant. Our findings suggest that eyebrow hair samples serve as a favorable tissue for germline genetic testing using next generation sequencing. Eyebrow hair follicle samples are easy to collect, making them a patient-friendly option for tissue procurement.

Our analysis also uncovers factors which should be considered in genetic germline testing, as they can offer early indications of the potential for a high percentage of donor-derived DNA in a patient. Firstly, we observed that sex matched transplantation is associated with an 8-percentage point higher donor derived DNA share. Secondly, the duration of time elapsed post-transplantation is negatively correlated with donor DNA shares. This underscores a dynamic nature of donor derived DNA shares over time and highlights the significance of considering the time elapsed since transplantation. Furthermore, we found that a better HLA-match and a matched related donor are associated with significantly lower DNA shares in our observed tissues. We thus conclude that these factors—sex-matching, time elapsed post-transplantation, HLA-matching, and donor relatedness—should be considered prior to genetic germline testing.

Finally, we compared the effectiveness of two different analysis methods: next-generation sequencing (NGS) and a short tandem repeat (STR) assay, in detecting donor derived DNA shares across varying percentages. NGS results indicated lower percentages of donor-derived DNA. The high sensitivity and resolution of NGS technology enables the detection of subtle genetic variations with higher precision, making it particularly suitable for detecting samples with low levels of donor derived DNA. On the other hand, our results indicated that STR analysis exhibits comparable quality to NGS while detecting on average higher percentages of donor derived DNA shares. Further research is warranted to elucidate the mechanisms underlying donor-derived DNA presence in several tissues, particularly concerning the dynamic nature of donor-derived DNA over time and in the case of reoccurring of the underlying disease.

## V. Conclusion

In conclusion, our study demonstrates the potential of using eyebrow hair follicles for accurate NGS analysis in germline genetic testing after stem cell transplantation due to its exceptionally low donor-derived DNA content. Factors signaling a higher donor-derived DNA share should be considered before germline genetic testing and include same-sex transplantation, a shorter time post-transplantation, low HLA match, and a non-related donor. NGS consistently outperforms STR analysis, offering greater sensitivity and precision for detecting low donor DNA percentages. To enhance the reliability and confidence of germline analysis, parallel SNP assays including donor derived DNA are recommended alongside NGS to rule out high donor DNA content. Figure 3 Error! Reference source not found. outlines a protocol for genetic germline testing following transplantation.

**FIGURE 3:**
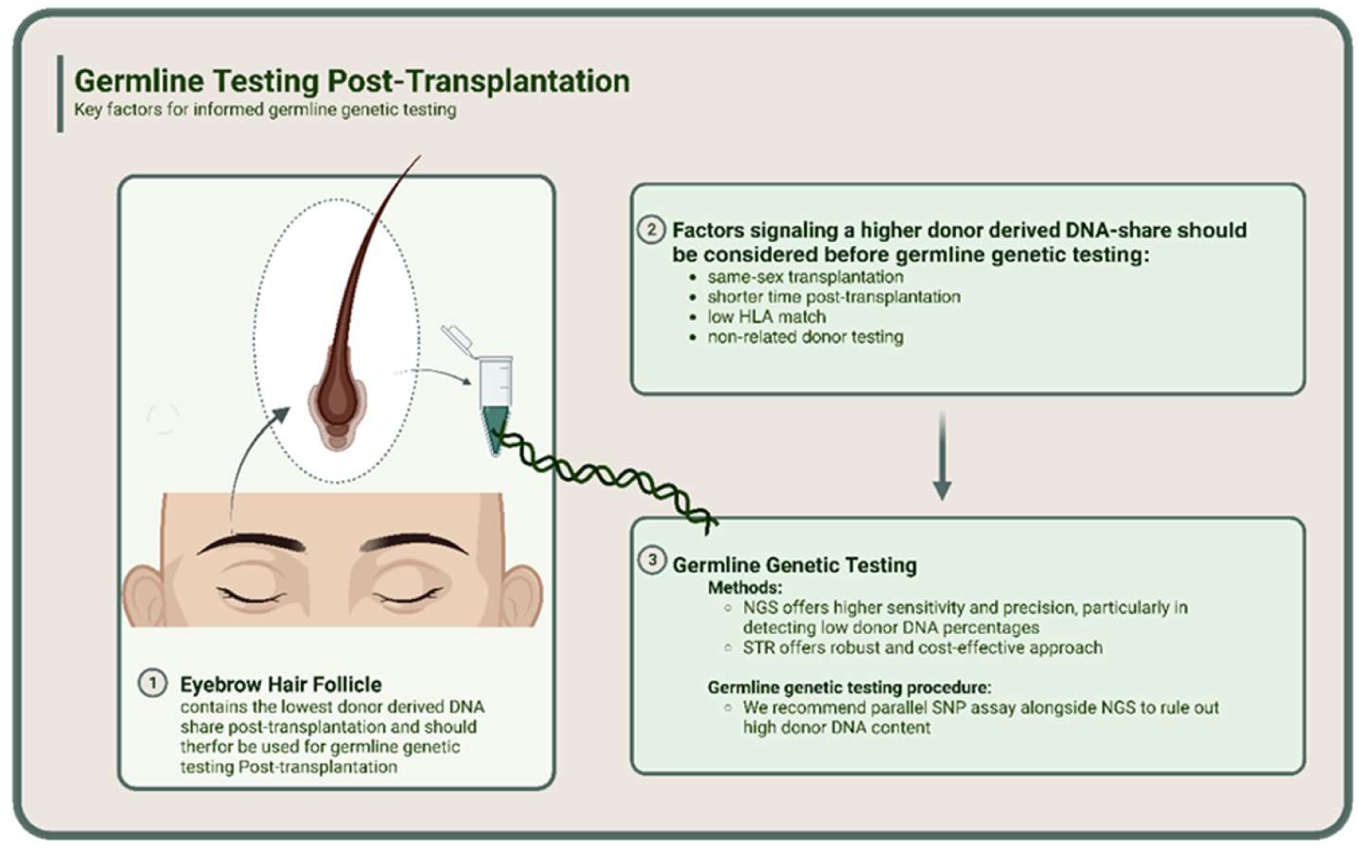
Protocol for germiline Testing post-transplantation.

Moving forward, research should prioritize the development of methodologies to distinguish between low-level mosaicism and donor-derived DNA in hair samples. Current NGS data analysis lacks definitive discrimination between these scenarios due to their low allele frequency. Advancements in sequencing technology and data analysis algorithms are essential to improve sensitivity and specificity in detecting these subtle genetic changes.

## VI. Data Availability Statement

The analyzed data supporting the findings of this study are included in the published article. Additional raw data are available from the corresponding author upon reasonable request.

## VII. Code Availability

The stata code used for analyzing the data in this study is available from the corresponding author upon reasonable request.

## VIII. Author Contribution Statement

All authors contributed significantly to the intellectual content of this manuscript and meet the ICMJE criteria for authorshttp. Specifically:

**Matthias Mertens** (shared first author) conducted the statistical analysis, played a key role in data interpretation and manuscript drafting.

**Mona Sadlo** (shared first author) conducted the wet lab methods, played a key role in data collection, and manuscript drafting.

**Julia Hentschel** (shared last author) conceived our study, played a key role in study design, contributed her expertise in wet lab methods, contributed to data interpretation, and provided critical revisions to the manuscript.

**Mareike Mertens** (shared last author) played a key role in study design, played a key role in data interpretation, contributed her expertise in human genetics, provided critical revisions to the manuscript, and served as the corresponding author.

All authors have approved the final version of the manuscript and agree to be accountable for all aspects of the work, ensuring that any questions related to accuracy or integrity are appropriately investigated and resolved. The corresponding author confirms full access to all data in the study and takes final responsibility for the decision to submit the manuscript for publication.

## IX. Ethical Approval

This study was conducted in accordance with the ethical standards of the institutional and/or national research committee and with the 1964 Helsinki declaration and its later amendments or comparable ethical standards. The protocol was approved by the Ethics Committee of the Medical Faculty of the University of Leipzig (186/21-ek). Informed consent was obtained from all subjects involved in the study or their legal guardians.

## XI. Figure Legends

Figure 1 illustrates the results of the dilution series using NGS, and STR, applying our code. Panel A displays results using NGS. The x-axis shows the actual DNA proportions of the eight diluted samples ranging from 2 % to 50 %. The y-axis displays the measured DNA proportions based on our NGS analysis. Boxplots represent the distribution of DNA proportions within each dilution across all markers. Median DNA proportions are marked by a white Panel B presents the results of the STR analysis using our code. The x-axis shows the actual DNA proportions of the eight diluted samples ranging from 2 % to 50 %.

Figure 2 shows boxplots for patient-specific median values of the results of the STR analysis using our code. On the x-axis, the actual DNA proportions of the eight diluted samples range from 2% to 50%, while the y-axis displays the measured DNA proportions based on based on our STR analysis and code. Boxplots represent the distribution of DNA proportions within each dilution across all markers. Median DNA proportions are marked by a white

Figure 3 outlines a protocol for genetic germline testing following transplantation. Figure created with BioRender.com.

## XII. Figures

**TABLE A.1.**
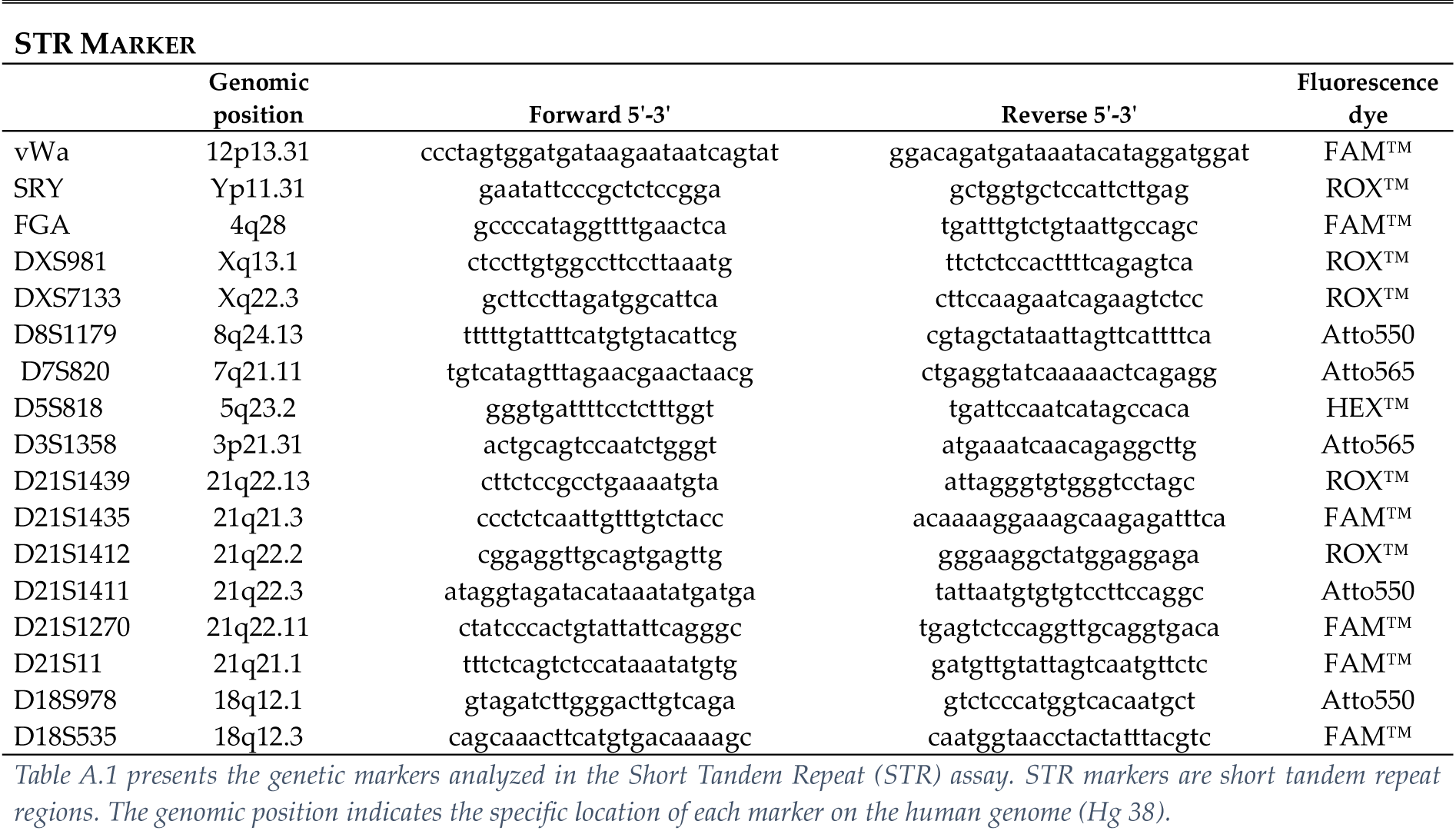
Genetic Markers analyzed in str Panels.

**TABLE A.2.**
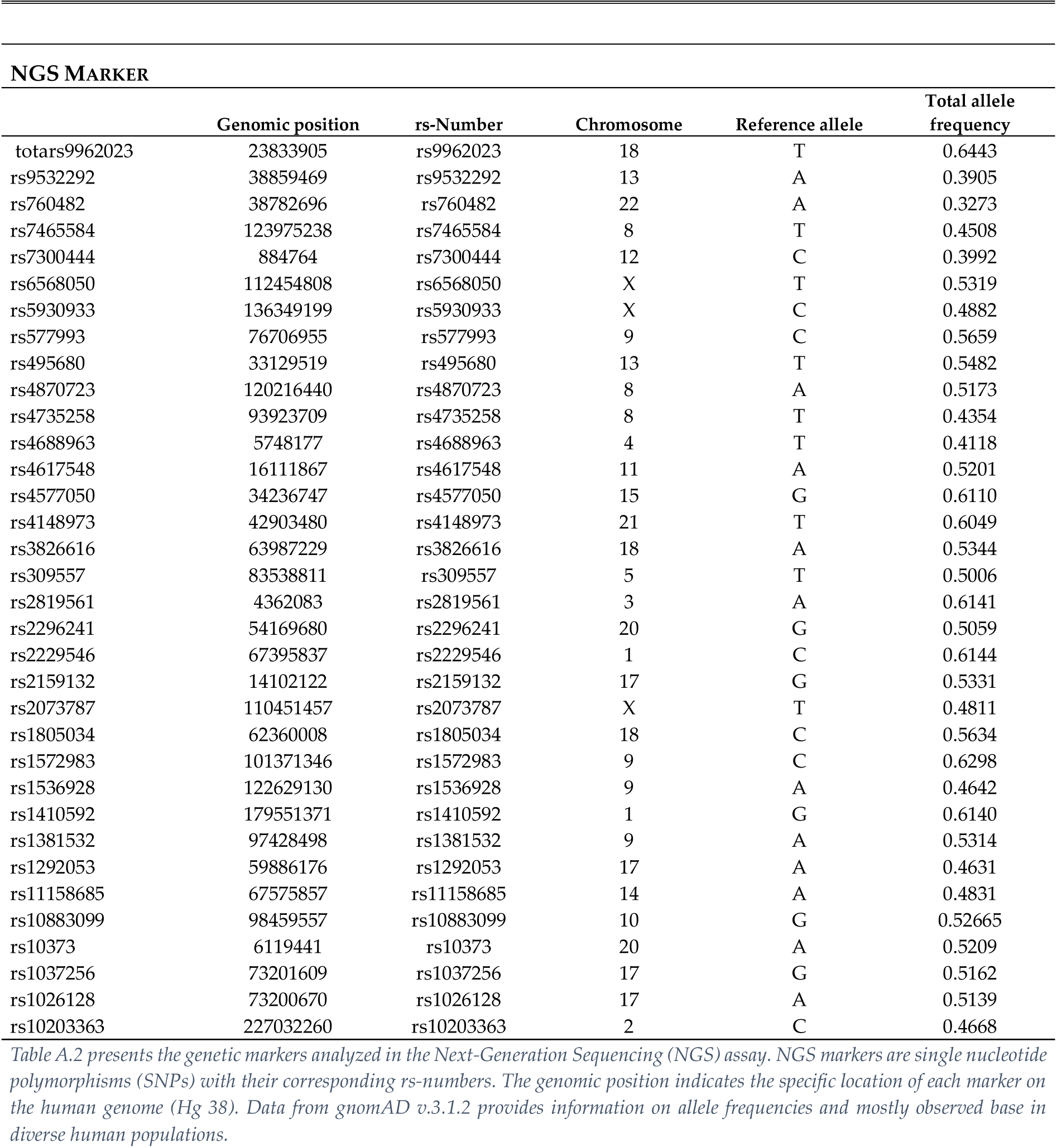
Genetic Markers analyzed in ngs Panels.

| GENETIC MARKERS ANALYZED IN NGS PANELS |  |  |  |  |  |
| --- | --- | --- | --- | --- | --- |
| NGS MARKER |  |  |  |  |  |
|  | Genomic position | rs-Number | Chromosome | Reference allele | Total allele frequency |
| totars9962023 | 23833905 | rs9962023 | 18 | T | 0.6443 |
| rs9532292 | 38859469 | rs9532292 | 13 | A | 0.3905 |
| rs760482 | 38782696 | rs760482 | 22 | A | 0.3273 |
| rs7465584 | 123975238 | rs7465584 | 8 | T | 0.4508 |
| rs7300444 | 884764 | rs7300444 | 12 | C | 0.3992 |
| rs6568050 | 112454808 | rs6568050 | X | T | 0.5319 |
| rs5930933 | 136349199 | rs5930933 | X | C | 0.4882 |
| rs577993 | 76706955 | rs577993 | 9 | C | 0.5659 |
| rs495680 | 33129519 | rs495680 | 13 | T | 0.5482 |
| rs4870723 | 120216440 | rs4870723 | 8 | A | 0.5173 |
| rs4735258 | 93923709 | rs4735258 | 8 | T | 0.4354 |
| rs4688963 | 5748177 | rs4688963 | 4 | T | 0.4118 |
| rs4617548 | 16111867 | rs4617548 | 11 | A | 0.5201 |
| rs4577050 | 34236747 | rs4577050 | 15 | G | 0.6110 |
| rs4148973 | 42903480 | rs4148973 | 21 | T | 0.6049 |
| rs3826616 | 63987229 | rs3826616 | 18 | A | 0.5344 |
| rs309557 | 83538811 | rs309557 | 5 | T | 0.5006 |
| rs2819561 | 4362083 | rs2819561 | 3 | A | 0.6141 |
| rs2296241 | 54169680 | rs2296241 | 20 | G | 0.5059 |
| rs2229546 | 67395837 | rs2229546 | 1 | C | 0.6144 |
| rs2159132 | 14102122 | rs2159132 | 17 | G | 0.5331 |
| rs2073787 | 110451457 | rs2073787 | X | T | 0.4811 |
| rs1805034 | 62360008 | rs1805034 | 18 | C | 0.5634 |
| rs1572983 | 101371346 | rs1572983 | 9 | C | 0.6298 |
| rs1536928 | 122629130 | rs1536928 | 9 | A | 0.4642 |
| rs1410592 | 179551371 | rs1410592 | 1 | G | 0.6140 |
| rs1381532 | 97428498 | rs1381532 | 9 | A | 0.5314 |
| rs1292053 | 59886176 | rs1292053 | 17 | A | 0.4631 |
| rs11158685 | 67575857 | rs11158685 | 14 | A | 0.4831 |
| rs10883099 | 98459557 | rs10883099 | 10 | G | 0.52665 |
| rs10373 | 6119441 | rs10373 | 20 | A | 0.5209 |
| rs1037256 | 73201609 | rs1037256 | 17 | G | 0.5162 |
| rs1026128 | 73200670 | rs1026128 | 17 | A | 0.5139 |
| rs10203363 | 227032260 | rs10203363 | 2 | C | 0.4668 |

**TABLE A.3.** Differences in donor-derived dna-shares across tissues.

| DIFFERENCES IN DONOR-DERIVED DNA-SHARES ACROSS TISSUES |  |  |  |  |  |  |
| --- | --- | --- | --- | --- | --- | --- |
| Variable | Mean<br>(1) | Median<br>(2) | SD<br>(3) | Max<br>(4) | Min<br>(5) | N<br>(6) |
| Median donor share per nail sample (NGS) | 0.19 | 0.08 | 0.24 | 0.89 | 0.00 | 45 |
| Median donor share per nail sample (STR) | 0.29 | 0.16 | 0.28 | 0.94 | 0.01 | 44 |
| Median donor share per buccal swab sample (NGS) | 0.25 | 0.20 | 0.18 | 0.87 | 0.00 | 45 |
| Median donor share per buccal swab sample (STR) | 0.29 | 0.24 | 0.19 | 0.88 | 0.03 | 45 |
| Median donor share per eyebrow follicle sample (NGS) | 0.01 | 0.01 | 0.01 | 0.04 | 0.00 | 28 |
| Median donor share per eyebrow follicle sample (STR) | 0.03 | 0.03 | 0.03 | 0.13 | 0.00 | 28 |

## Footnotes

ii We classify a marker as technical artefact or mosaic if the percentage of one base count lies between 60% and 90% or 10% and 40%. We set values between these intervals to 0%, 50%, and 100% to account for potential measurement error.

iii See Appendix Table A.3 for the associated statistics.

